# LRRK2 Suppresses Lysosome Degradative Activity in Macrophages and Microglia Through MiT-TFE Transcription Factor Inhibition

**DOI:** 10.1101/2022.12.17.520834

**Authors:** Narayana Yadavalli, Shawn M. Ferguson

## Abstract

Cells maintain optimal levels of lysosome degradative activity to protect against pathogens, clear waste and generate nutrients. Here we show that LRRK2, a protein that is tightly linked to Parkinson’s disease, negatively regulates lysosome degradative activity in macrophages and microglia via a transcriptional mechanism. Depletion of LRRK2 and inhibition of LRRK2 kinase activity enhanced lysosomal proteolytic activity and increased the expression of multiple lysosomal hydrolases. Conversely, the kinase hyperactive LRRK2 G2019S Parkinson’s disease mutant suppressed lysosomal degradative activity and gene expression. We identified MiT-TFE transcription factors (TFE3, TFEB and MITF) as mediators of LRRK2-dependent control of lysosomal gene expression. LRRK2 negatively regulated the abundance and nuclear localization of these transcription factors and their depletion prevented LRRK2-dependent changes in lysosome protein levels. These discoveries define a role for LRRK2 in controlling lysosome degradative activity and support a model wherein LRRK2 hyperactivity may increase Parkinson’s disease risk by suppressing lysosome degradative activity.

**Significance Statement:** This study defines a homeostatic mechanism that allows macrophages and microglia to match the degradative activity of their lysosomes to ongoing changes in cellular demand. It shows that the leucine rich repeat kinase 2 (LRRK2) protein suppresses lysosome degradative activity by inhibiting the expression and nuclear localization of the MiT-TFE family of transcription factors that control the expression of multiple genes that encode lysosome proteins. It further demonstrates that a Parkinson’s disease mutation that hyperactivates LRRK2 kinase activity limits the degradative activity of lysosomes more strongly. These findings support a model wherein LRRK2 protects cells from excessive lysosome degradative activity and suggest that overactivation of this pathway may increase Parkinson’s disease risk by limiting the degradative activity of lysosomes.

## Introduction

Lysosomes serve important functions as sites of pathogen defense, macromolecule degradation, intracellular nutrient release and as signaling platforms that coordinate cellular responses to nutrient and growth factor availability^1, 2^. Cells must therefore control the activity of their lysosomes to match ongoing changes in cellular demand. Major advances in understanding the signaling and homeostatic control of lysosomes were the discoveries that proteins of the MiT-TFE family of transcription factors stimulate the expression of multiple genes that encode lysosomal proteins and are in turn regulated by lysosomes and the nutrients that they release^3–12^. Central to this process are tightly regulated interactions of the MiT-TFE proteins with Rag GTPases and the mechanistic target of rapamycin complex 1 (mTORC1) that occur on the surface of lysosomes^6–9, 11–13^.

In addition to the established role for mTORC1 and MiT-TFE transcription factors in coordinating lysosome homeostasis, recent studies have proposed roles for leucine rich repeat kinase 2 (LRRK2) in supporting the health of lysosomes. LRRK2 is recruited to the surface of lysosomes in specialized cell types including macrophages and astrocytes in response to stimuli that stress and/or damage lysosome membranes^14–17^. This recruitment of LRRK2 to lysosome membranes has been proposed to promote lysosome membrane repair^14, 15, 18^. LRRK2-dependent regulation of lysosome protease activity has also been observed in macrophages^19^. Broader functional relevance of LRRK2 for lysosomes is supported by observations of additional lysosome-related phenotypes following LRRK2 inhibition or knockout in a range of cells and tissues^20–26^. This includes both changes to lysosome morphology as well as increases in the abundance of multiple lysosome proteins via mechanisms that remain poorly understood^20–26^.

In addition to these emerging cell biological insights about LRRK2 and lysosomes, mutations in the human *LRRK2* gene cause an autosomal dominant form of Parkinson’s disease and additional LRRK2 variants contribute to risk for sporadic Parkinson’s disease^27–30^. Human genetics studies have furthermore linked LRRK2 to risk for pathogen infections and inflammatory diseases^31–33^. These connections between LRRK2 and human diseases raise questions about the normal functions of LRRK2 and the mechanisms whereby excessive or aberrant LRRK2 promotes disease. Considerable attention has focused on the kinase activity of LRRK2 due to the fact that multiple Parkinson’s disease mutations increase this activity^34, 35^. This has motivated the development of LRRK2 inhibitors as a putative Parkinson’s disease therapy^36^.

Given the major impact of Parkinson’s disease on midbrain dopaminergic neurons, significant attention has focused on neuronal roles for LRRK2^37–41^. However, LRRK2 is also expressed in diverse non-neuronal cell types. In particular, LRRK2 expression levels are notable in cells of the myeloid lineage including macrophages and microglia^17, 19, 31, 42–44^. The potential Parkinson’s disease importance of microglial LRRK2 is supported by a recent analysis of human genetics data that identified a non-coding LRRK2 variant that confers Parkinson’s disease risk via regulation of LRRK2 expression in microglia^45^. Functions for LRRK2 in microglia and related myeloid cells such as macrophages are also supported by studies that have described phenotypes arising from LRRK2 perturbations in such cells^14, 16, 17, 19, 31, 46–49^.

To investigate the relationship between LRRK2 and lysosomes in macrophages and microglia, we took advantage of LRRK2 mutant mouse models as well as the recent development of protocols for the differentiation of macrophages and microglia from human iPSCs to test the impact of LRRK2 genetic and pharmacological perturbations on lysosomal degradative activity in these specialized cell types. Our results revealed an inverse relationship between LRRK2 kinase activity and the expression of genes encoding lysosome proteins. In search of a mechanism to explain LRRK2-dependent changes in lysosome gene expression, we identified a major role for LRRK2 in suppressing the abundance and nuclear localization of MiT-TFE transcription factors and established a requirement for them in mediating LRRK2-dependent changes in the abundance of key lysosome proteins. Collectively, this data supports a model wherein LRRK2 suppresses the degradative activity of lysosomes in macrophages and microglia by negatively regulating the MiT-TFE transcription factors. Mutations or other factors that activate LRRK2 kinase activity may therefore increase Parkinson’s disease risk through lysosome inhibition.

## Results

### LRRK2 suppresses lysosome degradative activity in human and mouse macrophages

To investigate the relationship between LRRK2 and macrophage lysosome degradative activity, we took advantage of an established protocol to differentiate control and LRRK2 KO human iPSCs into macrophages^50, 51^. Both tubular and vesicular lysosomes were robustly labeled in these macrophages following incubation with Alexa488-BSA and DQ-BSA which identify lysosomes and report their proteolytic activity respectively^52^ (Fig. 1A). This combination of probes allows the measurement of lysosomal proteolytic activity while controlling for potential changes in endocytic uptake. Inhibition of LRRK2 kinase activity in control macrophages with two different drugs (MLi-2 and LRRK2-IN-1) resulted in an increase in the DQ-BSA signal without affecting overall endocytic uptake as measured by the Alexa488-BSA signal (Fig. 1B and C). A similar increase in the DQ-BSA signal was observed in LRRK2 KO macrophages and this was not further enhanced by either of the LRRK2 inhibitors (Fig. 1B and C). The lack of any additional effect of these inhibitors on the LRRK2 KO cells argues against off target effects of these drugs as drivers of changes in lysosome proteolytic activity. In time course experiments, differences in the DQ-BSA signal in response to LRRK2 inhibition with MLi-2 were detectable by 3 hours and reached a maximum after 12-24 hours of treatment (Fig. S1A).

**Figure 1.**
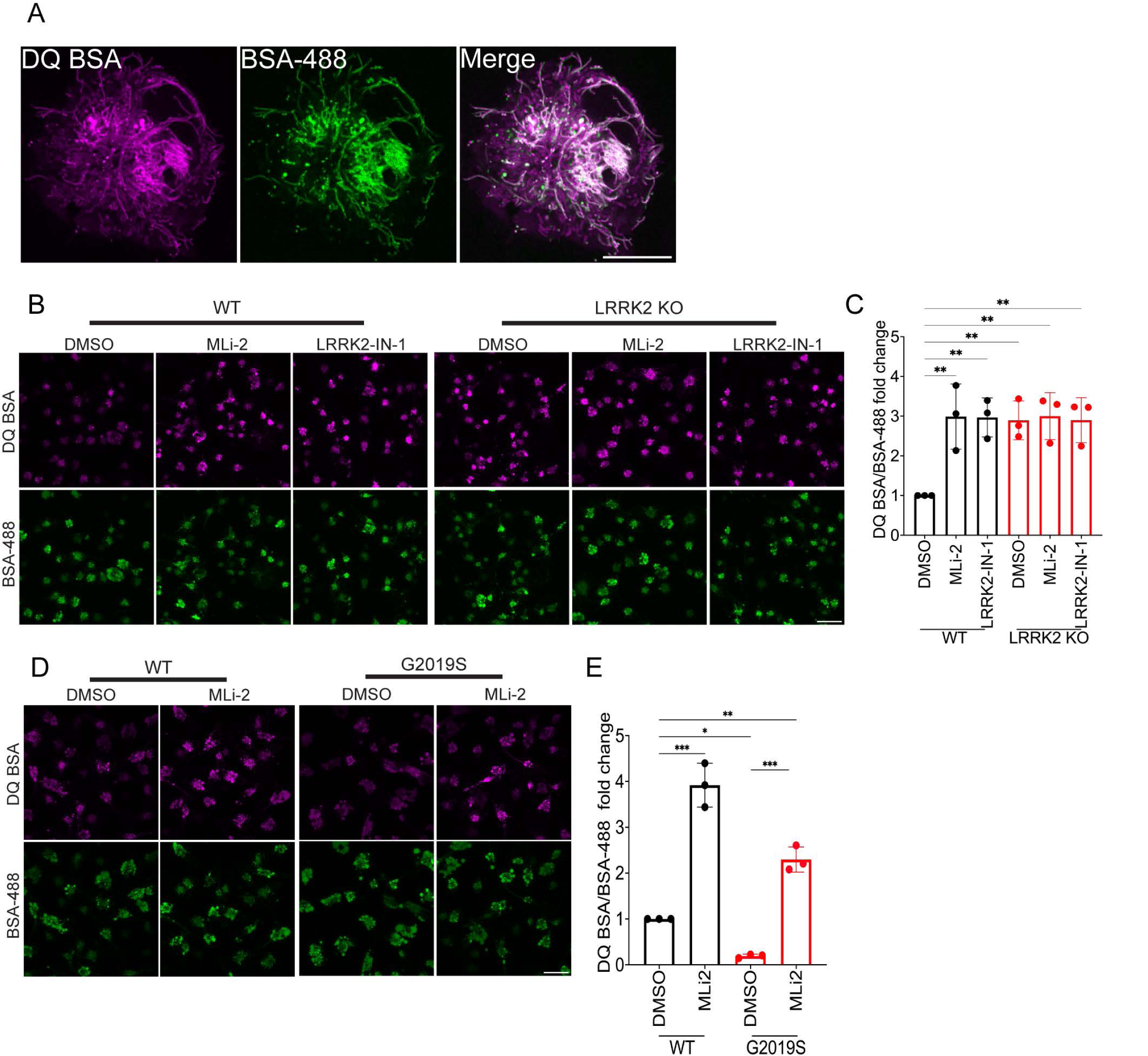
LRRK2 inhibits lysosomal protease activity in human iPSC-derived macrophages. **(A)** Airyscan live cell confocal micrographs showing lysosome labeling in human iPSC-derived macrophages following labeling with DQ-BSA and Alexa488-BSA. Scale bar, 5 µm. **(B)** Confocal micrographs showing the DQ-BSA and Alexa488-BSA fluorescence in WT and LRRK2 KO macrophages treated with 0.1% DMSO (vehicle), 50 nM MLi-2 and 250 nM LRRK2-IN-1 respectively for 3 hours. Scale bar, 10 µm. **(C)** The mean fluorescence intensity of DQ-BSA was normalized to BSA-488 and plotted relative to the DMSO control. The data was collected from 3 independent experiments with 60-80 cells per experiment. **(D)** Confocal micrographs showing the DQ-BSA and Alexa488-BSA fluorescence in WT and LRRK2 G2019S mutant macrophages. Scale bar, 10 µm. **(E)** Bar graph showing the quantification of DQ-BSA/BSA 488 fluorescence in WT and LRRK2 G2019S macrophages. The data was collected from 3 independent experiments with 60-80 cells per experiment. Error bars show mean ± SEM, one-way ANOVA, * p<0.05, ** p< 0.01, ***p< 0.001.

To test the generalizability of these results, we performed additional experiments in primary cultures of both wildtype and *Lrrk2* KO mouse bone marrow derived macrophages (BMDMs). In each case, the KO of LRRK2 or inhibition of its kinase activity resulted in an increase in lysosomal protease activity as measured by the DQ-BSA assay without any concomitant change in the total labeling of lysosomes by Alexa488-Dextran (Fig. S1 B and C).

### LRRK2 G2019S Parkinson’s disease mutant suppresses macrophage lysosome proteolytic activity

Based on the increased lysosome proteolytic activity following genetic and pharmacological inhibition of LRRK2, we predicted that the gain-of-function LRRK2 G2019S Parkinson’s disease mutant which has increased kinase activity would have the opposite effect. Indeed, macrophages derived from human iPSCs with a knockin of the G2019S mutation had reduced lysosomal protease activity and this was rescued by LRRK2 inhibition (Fig. 1D and E). The same was true in BMDMs from G2019S knockin mice (Fig. S1D and E).

### LRRK2 negatively regulates the abundance of multiple lysosomal proteins

We next performed immunoblotting experiments on human iPSC-derived macrophages to determine whether the LRRK2-dependent changes in lysosome proteolytic activity were associated with changes in the abundance of lysosome proteases. These experiments revealed increases in the levels of cathepsins B, C, D and L in response to either LRRK2 KO or incubation with LRRK2 inhibitors and the effects of the KO and the inhibitors were not additive (Fig. 2A-E). These effects were not limited to lysosomal proteases as levels of glucocerebrosidase (GCase/GBA), a Parkinson’s linked lysosomal enzyme involved in glycosphingolipid metabolism, also increased in response to LRRK2 inhibition (Fig. 2A and F)^53^. This effect also extended to LAMP1, a major membrane glycoprotein of lysosomes (Fig. 2A and G). Similar changes were observed in WT versus *Lrrk2* KO mouse bone marrow derived macrophages (Fig. S2A-F). Time course experiments detected lysosome protein changes as early as 3 hours after initiation of MLi-2 treatment and these changes increased through 12-24 hours of treatment (Fig. 2H-J).

**Figure 2:**
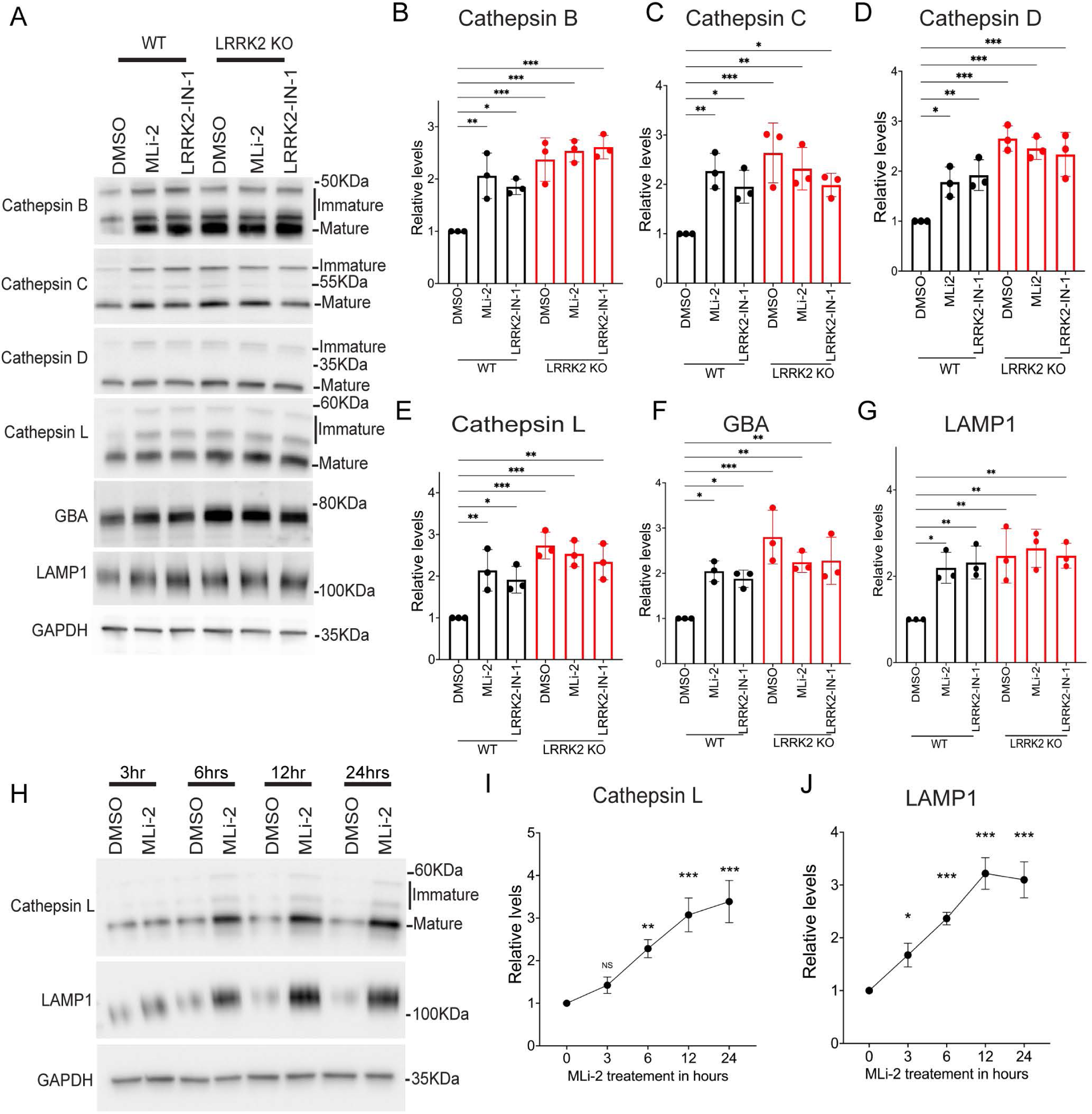
LRRK2 negatively regulates the levels of multiple lysosomal proteins in human iPSC-derived macrophages. **(A)** Immunoblot analysis of WT and LRRK2 KO macrophages treated with 0.1% DMSO (vehicle), 50 nM MLi-2 or 250 nM LRRK2-IN-1 for 6 hours. **(B-G)** Quantification of immunoblots in A. For samples where multiple bands reflect immature and mature cathepsin proteins, the quantification reflects the total of all bands. **(H)** Immunoblots showing the time course analysis of Cathepsin L and LAMP1 in response to 50 nM MLi-2. **(I-J)** Quantification of immunoblots from panel G. For all experiments, quantification was performed on data collected from 3 independent experiments and plotted in relative to DMSO control cells. Error bars show mean ± SEM, one-way ANOVA, * p<0.05, ** p< 0.01, ***p< 0.001.

Meanwhile, knockin of the *LRRK2* G2019S mutation in both human iPSC-derived macrophages and mouse bone marrow derived macrophages resulted in reduced levels of lysosome hydrolases and LAMP1 and this was reversed by LRRK2 inhibition (Fig. 3A-G and S3A-D).

**Figure 3:**
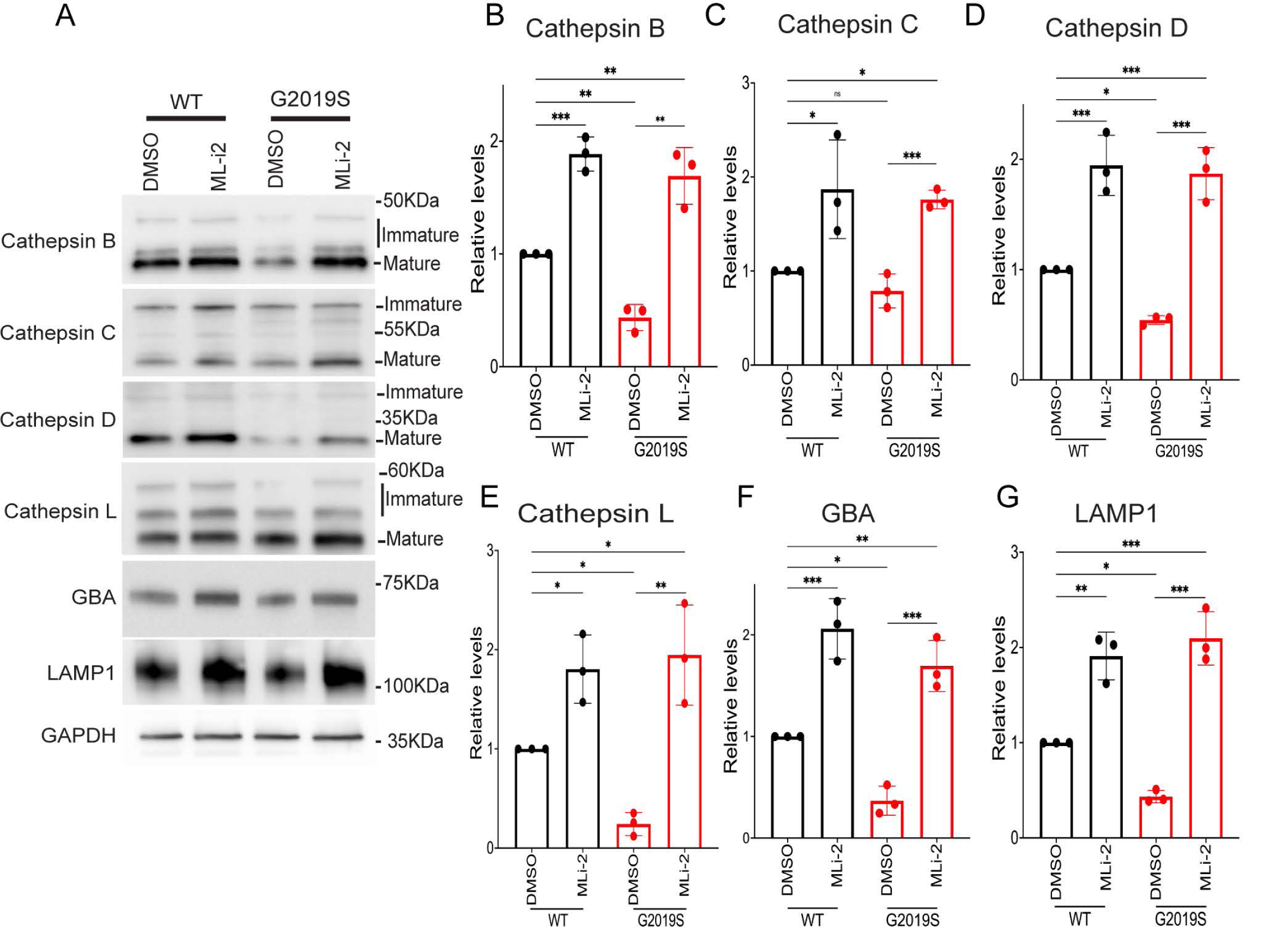
LRRK2 G2019S autosomal dominant Parkinson’s disease mutation decreases levels of lysosomal proteins in human iPSC-derived macrophages. **(A)** Immunoblot analysis of WT and LRRK2 G2019S knockin macrophages treated with 0.1% DMSO (vehicle), 50 nM MLi-2 or 250 nM LRRK2-IN-1 for 6 hours. **(B-G)** Quantification of immunoblots in A. For samples where multiple bands reflect immature and mature cathepsin proteins, the quantification reflects the total of all bands. Data was collected from 3 independent experiments and plotted in relative to DMSO control cells. Error bars show mean ± SEM, one-way ANOVA, * p<0.05, ** p< 0.01, ***p< 0.001.

### LRRK2 suppresses the expression of multiple mRNAs that encode lysosome proteins

Newly made cathepsins are delivered to lysosomes in “pro” forms and then undergo activation via proteolytic processing. As a result, they exhibit band patterns on immunoblots that reflect this coupling between trafficking and processing. LRRK2 genetic and pharmacological perturbations increased all forms of these cathepsins and not just the abundance of the mature proteins (Figure 2, 3, S2-S4). As this suggested an increase in cathepsin synthesis, we next performed qRT-PCR and observed that LRRK2 KO and LRRK2 inhibition both resulted in an increase in the expression of multiple transcripts that encode lysosomal proteins (Fig. 4A-D). Meanwhile, these transcripts were down-regulated in G2019S knockin macrophages and this was rescued by LRRK2 inhibition (Fig. 4E-H). Time course experiments revealed that these mRNA expression changes in response to LRRK2 inhibition were detectable as early as 3 hours and reached a maximum by 12 hours (Fig. 4 I+J).

**Figure 4:**
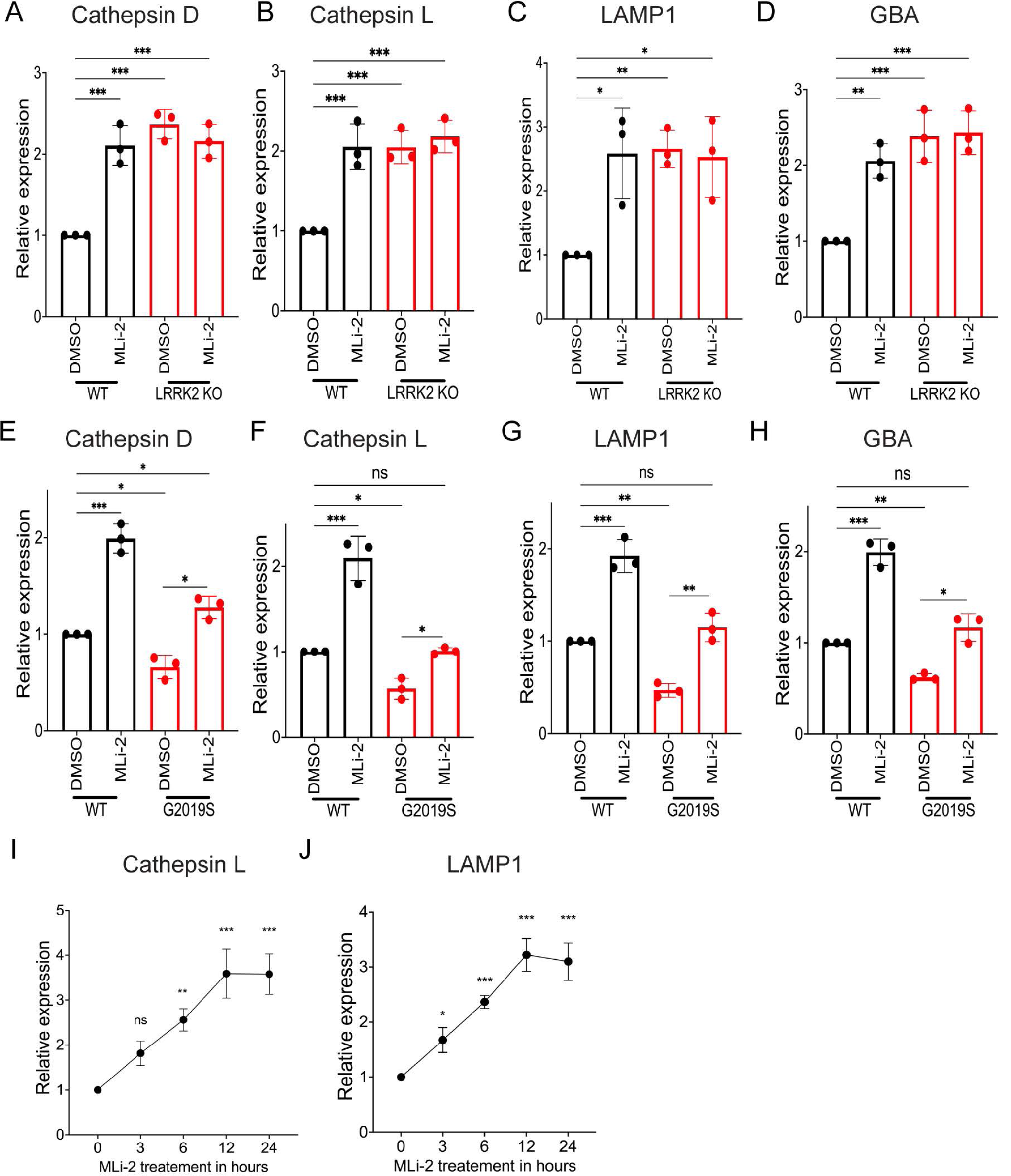
LRRK2 negatively regulates mRNA expression for multiple lysosome-related genes in human iPSC-derived macrophages. **(A-D)** qRT-PCR analysis of lysosomal transcripts (Cathepsin D, Cathepsin L, GBA and LAMP1) in WT and LRRK2 KO macrophages treated with 0.1% DMSO or 50 nM MLi-2 for 6 hours. Data was collected from 3 independent experiments, normalized to GAPDH and presented relative to DMSO control. **(E-H)** qRT-PCR analysis of lysosomal transcripts in WT and LRRK2 G2019S macrophages treated with 0.1% DMSO and 50nM MLi-2 for 6 hours. **(I and J)** qRT-PCR analysis of Cathepsin L and LAMP1 transcripts at the indicated timepoints of 50 nM MLi-2 treatment. Data was collected from 3 independent experiments, normalized to GAPDH and presented relative to DMSO control. Error bars show mean ± SEM, one-way ANOVA, * p<0.05, ** p< 0.01, ***p< 0.001.

### LRRK2 negatively regulates MiT-TFE transcription factors

In search of an explanation for how LRRK2 regulates the expression of multiple genes that encode lysosome proteins, we next focused on the MiT-TFE family of transcription factors as they coordinate the expression of genes encoding lysosome proteins by binding to a response element (CLEAR motif) that is found in their promoters^4, 54–56^. Past studies have shown that these transcription factors are highly regulated at the level of their nuclear versus cytoplasmic distribution^3, 9, 13, 56^. TFE3 stands out for having high expression in macrophages and also in microglia within the mouse brain^57, 58^. We therefore tested the effect of LRRK2 perturbations on TFE3 subcellular distribution. Endogenously expressed TFE3 was mostly excluded from the nucleus of control iPSC-derived macrophages but it became concentrated in the nucleus following LRRK2 inhibition (Fig. 5A and B). Furthermore, TFE3 was constitutively enriched in the nucleus in LRRK2 KO cells (Fig. 5A and B). This LRRK2-dependent negative regulation of TFE3 nuclear localization was also observed in mouse BMDMs in response to LRRK2 inhibition and KO as assessed both by confocal immunofluorescence and sub cellular fractionation assays (Fig. S5A-F). Meanwhile, both human and mouse macrophages with a knockin of the LRRK2 G2019S mutation had weak overall TFE3 immunofluorescence (Fig. 5C and D; S5G-H). This signal became stronger overall and concentrated in the nucleus in response to inhibition of LRRK2 kinase activity (Fig. 5C and D; S5G and H). Confocal immunofluorescence analysis of the localization of the endogenous TFEB protein in mouse macrophages also revealed that its nuclear localization is negatively regulated by LRRK2 in a manner that parallels what we observed for TFE3 following LRRK2 inhibition, LRRK2 KO and knockin of the G2019S mutation (Fig. S6 A-D). In contrast, in iPSC-derived cortical neurons, LRRK2 inhibition did not result in nuclear localization of TFE3 (Fig. S7). This result indicates that there is at least some cell type specificity for effects of LRRK2 inhibition on TFE3 subcellular localization.

**Figure 5:**
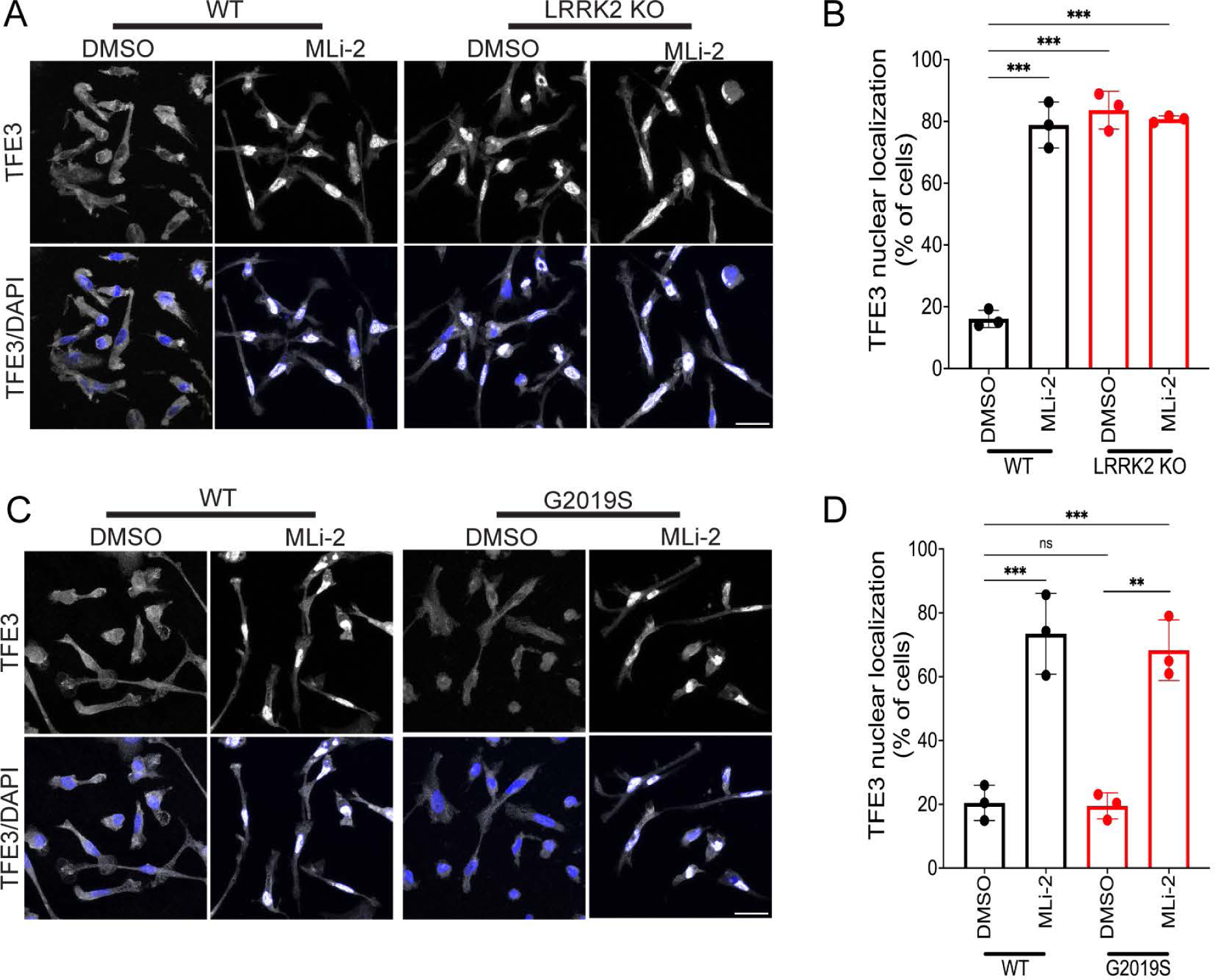
LRRK2 suppresses nuclear translocation of TFE3 in human iPSC-derived macrophages. **(A)** Immunofluorescence confocal micrographs showing the sub cellular localization of TFE3 in WT and LRRK2 KO macrophages treated with 0.1% DMSO and 50nM MLi-2 for 6 hours. **(B)** Quantification of TFE3 nuclear localization under the indicated conditions where cells were scored for having nuclear>cytoplasmic TFE3 signal. Data was collected from 3 independent experiments, approximately 60-70 cells were analyzed per experiment. **(C)** Immunofluorescence, confocal micrographs showing the sub cellular localization of TFE3 in WT and LRRK2 G2019 mutant macrophages after treatment with 0.1% DMSO and 50nM MLi-2 for 6 hours. **(D)** Quantification of % cells with nuclear > cytoplasmic TFE3. Data was collected from 3 independent experiments, approximately 50-70 cells per experiment.

In support of a causal role for MiT-TFE proteins in the LRRK2-dependent changes in lysosomal gene expression, LRRK2 inhibition no longer resulted in an increase in lysosome proteins following siRNA-mediated knockdown of TFE3 (Fig. 6A-E). As TFE3 overlaps with TFEB and MITF in controlling lysosome gene expression, we also examined their levels and found that they were also reduced in response to the TFE3 siRNA (Fig. 6F-H)^3^.

**Figure 6:**
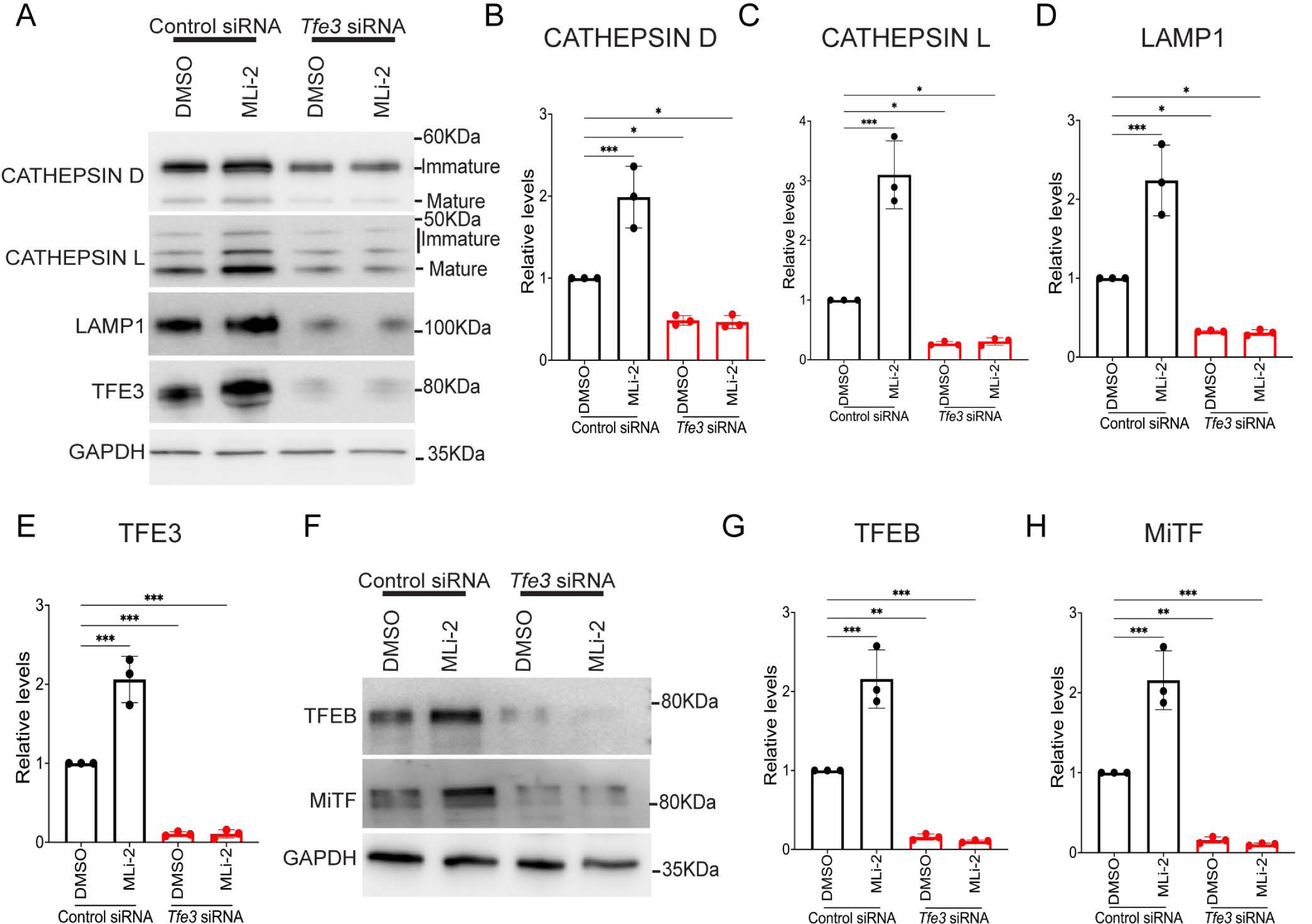
TFE3 depletion results in loss of LRRK2-dependent regulation of TFEB and MiTF. **(A)** Immunoblots showing the impact of control and Tfe3 siRNAs on the levels of the indicated proteins in cells that were treated with 0.1% DMSO or 50nM MLi-2 for 6 hours. **(B-E)** Quantification of the abundance of the indicated proteins from 3 independent experiments. **(F)** Immunoblots for TFEB, MITF and GAPDH from mouse macrophages treated with control and TFE3 siRNAs. **(G and H)** Quantification of immunoblots in panel F. Error bars show mean ± SEM, one-way ANOVA, * p<0.05, ** p< 0.01, ***p< 0.001.

The loss of TFEB and MITF upon TFE3 silencing suggests a positive feedback mechanism wherein TFE3 promotes the expression of itself as well as TFEB and MITF in macrophages. Consistent with this hypothesis, TFE3, TFEB and MITF protein levels all increased in response to LRRK2 inhibition with MLi-2 and this was abolished by TFE3 knockdown (Fig. 6A, E-H; Fig. 7A-D). The effect of LRRK2 on TFE3, TFEB and MITF abundance was also seen following treatment with LRRK2-IN-1 and in response to LRRK2 KO in human and mouse macrophages (Fig. 7A-D; S8A-D). Meanwhile, TFE3, TFEB and MITF protein levels were lower in LRRK2 G2019S human and mouse macrophages and this was rescued by treatment with MLi-2 (Fig. 7E-H; S8E-J). Consistent with a transcriptional mechanism for the LRRK2-dependent regulation of TFE3 expression, qRT-PCR assays revealed reciprocal effects of LRRK2 inhibition and the G2019S mutation on TFE3 transcript abundance (Fig. 7I). These experiments establish that in addition to controlling the nuclear localization of TFE3 and TFEB, LRRK2 also controls their abundance. Although MITF abundance shared similar regulation to TFE3 and TFEB, MITF subcellular localization was not investigated as its levels of expression in macrophages did not allow for robust detection in our imaging assays.

**Figure 7:**
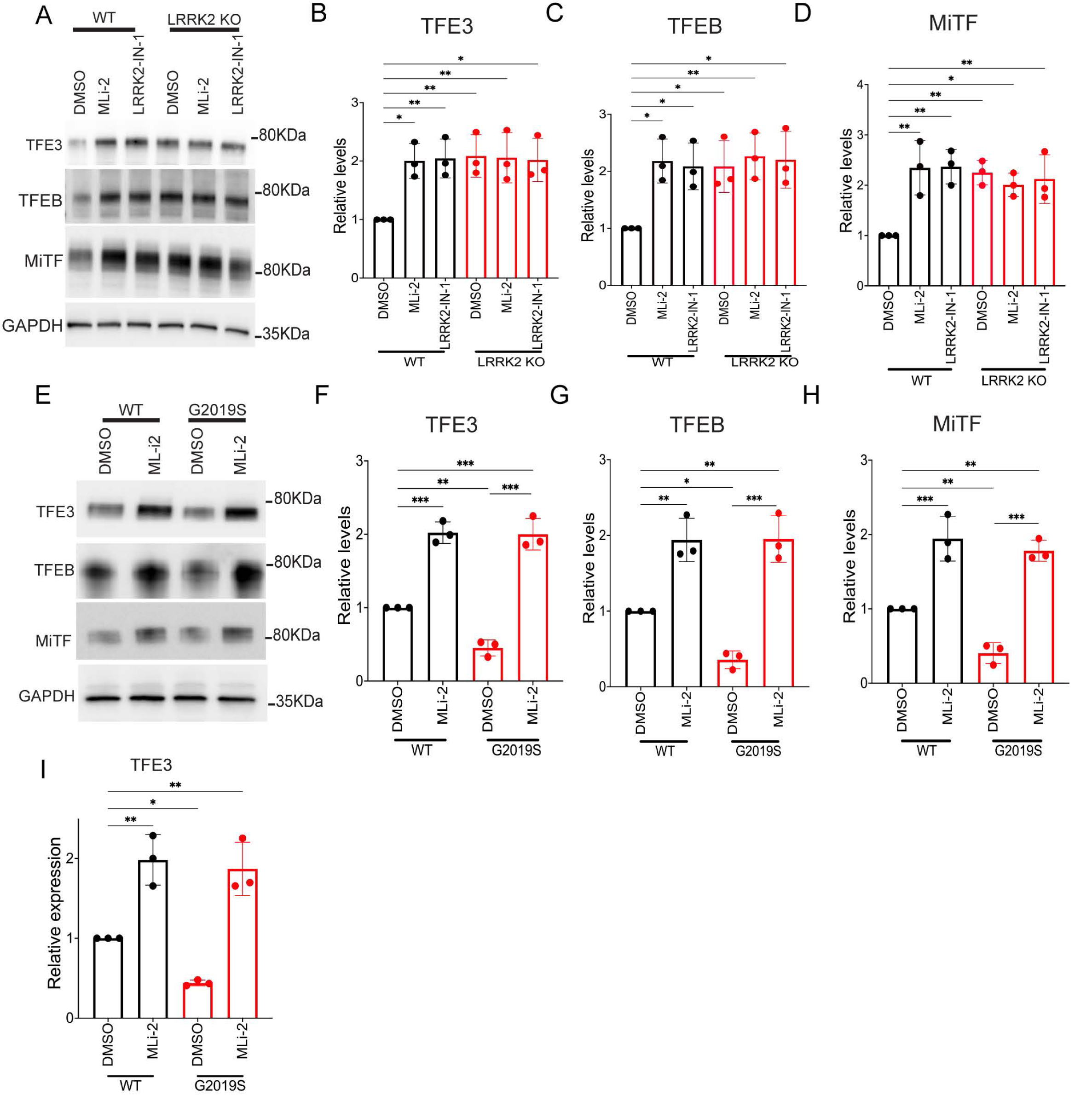
LRRK2 negatively regulates the protein levels of TFE3, TFEB and MITF in human iPSC-derived macrophages. **(A)** Immunoblots showing the levels of TFE3, TFEB and MITF proteins in WT and LRRK2 macrophages cells treated with 0.1% DMSO, 50nM MLi-2 or 250nM LRRK2-in-1 for 6 hours. **(B-D)** Quantification of immunoblots in A. Data was collected from 3 independent experiments and plotted relative to DMSO-treated WT cells. **(E)** Immunoblots showing the levels of TFE3, TFEB and MITF in LRRK2 G2019S mutant macrophages treated with 0.1% DMSO or 50nM MLi-2 for 6 hours. **(F-H)** Quantification of immunoblots in E. Data was collected from 3 independent experiments and plotted relative to DMSO-treated WT cells. **(I)** qRT-PCR analysis of TFE3 expression in WT versus LRRK2 G2019S mutant macrophages treated with 0.1% DMSO or 50nM MLi-2 for 6 hours. Data was collected from 3 independent experiments and plotted relative to DMSO-treated WT cells. Error bars show mean ± SEM, one-way ANOVA, * p<0.05, ** p< 0.01, ***p< 0.001.

### LRRK2-dependent regulation of microglial lysosomes

Given the impact of LRRK2 mutations in Parkinson’s disease, the close functional relationship between macrophages and microglia and recent genetic insights into the potential importance of microglial LRRK2 in Parkinson’s disease^17, 45^, we next differentiated human iPSCs (control, LRRK2 KO and G2019S) into microglia via an established protocol^59^. Similar to what we observed for human and mouse macrophages, microglia lysosome proteolytic activity increased in response to LRRK2 inhibition and LRRK2 KO and decreased in cells with the LRRK2 G2019S mutation (Fig. 8A-B). These observations were paralleled by increased lysosome hydrolase, LAMP1 and MiT-TFE protein abundance in response to LRRK2 inhibition and KO and their decreased abundance in the G2019S microglia (Fig. 8C-L).

**Figure 8:**
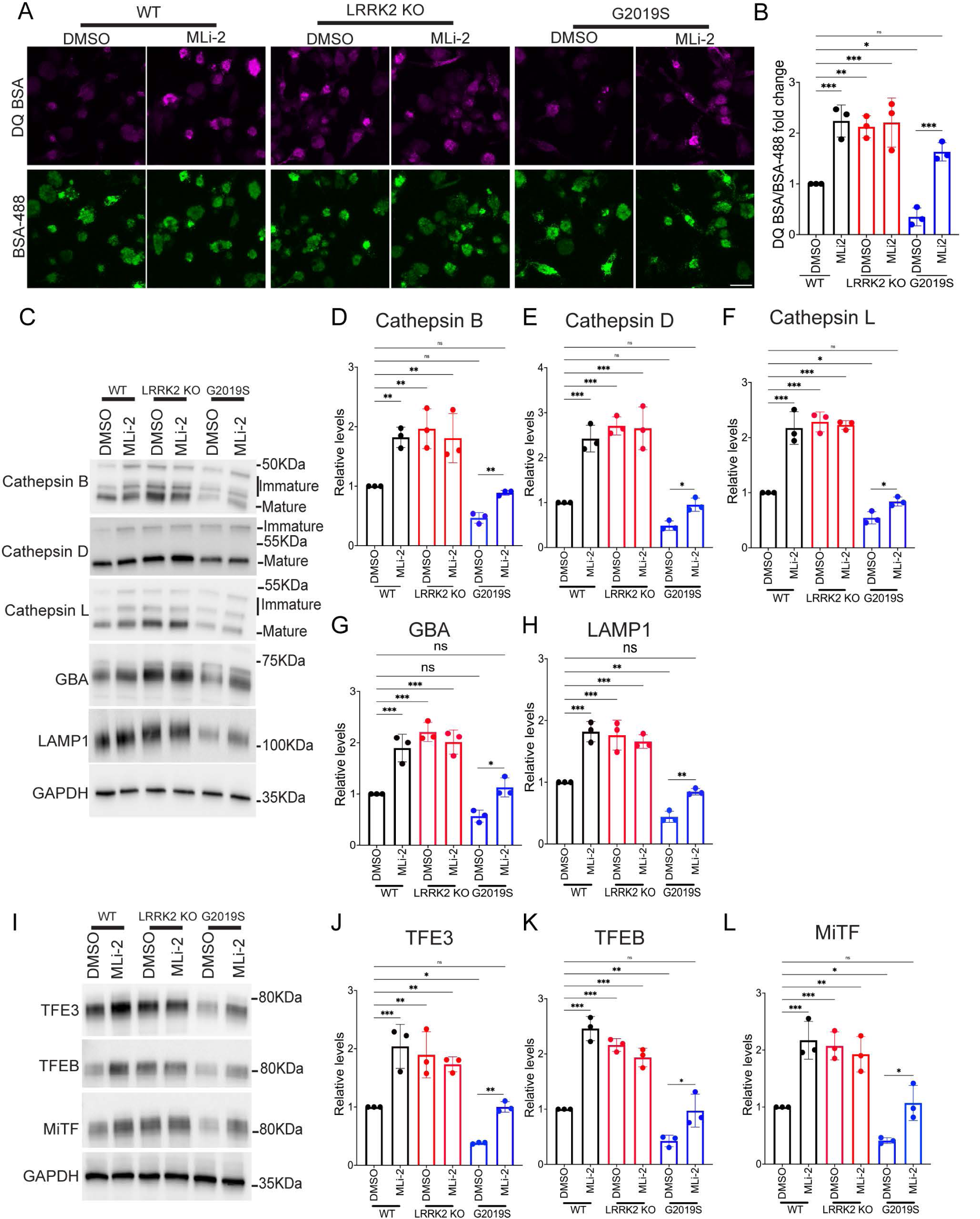
Regulation of human iPSC-derived microglial lysosomes by LRRK2. **(A)** Confocal micrographs showing DQ-BSA and Alexa488-BSA fluorescence in WT, LRRK2 KO and LRRK2 G2019S mutant macrophages treated with 0.1% DMSO or 50nM MLi2 for 3 hours. Scale bar, 10 µm. **(B)** Quantification of DQ-BSA/Alexa488-BSA fluorescence in WT, LRRK2 KO and LRRK2 G2019S mutant macrophages. The mean fluorescence intensity of DQ-BSA was normalized to BSA-488 and plotted relative to the DMSO control. The data was collected from 3 independent experiments with 60-80 cells per experiment. **(C)** Immunoblot analysis of WT, LRRK2 KO and LRRK2 G2019S mutant microglia treated with 0.1% DMSO, 50nM MLi-2 or 250 nM LRRK2-IN-1 for 6 hours. **(D-H)** Quantification of immunoblots for the indicated proteins from panel C. For the cathepsins, all bands (reflecting immature and mature proteins) were measured. Data was collected from 3 independent experiments, normalized to GAPDH abundance and plotted relative to DMSO-treated WT cells. **(I)** Immunoblots showing the levels of TFE3, TFEB and MITF proteins in WT, LRRK2 KO and G2019S knockin microglia treated for 6 hours with 0.1% DMSO or 50 nM MLi2. **(J-L)** Quantification of immunoblots from panel I. Data was collected from 3 independent experiments and plotted in relative to DMSO-treated WT cells. Error bars represent mean ± SEM; one-way ANOVA, * p<0.05, ** p< 0.01, ***p< 0.001.

## Discussion

Our results, based on the analysis of endogenously expressed proteins in human and mouse cells, demonstrate that LRRK2 suppresses lysosome degradative activity in macrophages and microglia by regulating the abundance and nuclear localization of MiT-TFE transcription factors. Given that lysosome degradative activity is generally thought of as a positive process that converts waste into nutrients, questions arise concerning why it would be advantageous to have a LRRK2-based mechanism for suppressing this activity. Building on the recently proposed roles for LRRK2 activation on the surface of damaged lysosomes^14, 15^, we speculate that LRRK2-dependent inhibition of TFE3, TFEB and MITF may act as a homeostatic mechanism to protect cells from leakage of lysosomal hydrolases into the cytoplasm. Such a brake might be particularly relevant in macrophages and microglia as these cells require highly degradative lysosomes to meet the demands imposed by their robust phagocytic activity but this also makes them vulnerable to perturbations to lysosome integrity that would result in the leakage of hydrolases into the cytoplasm^14, 15, 60^. Thus, when macrophage/microglia lysosome membranes are damaged, LRRK2 could act in parallel to both support local lysosome membrane repair processes and to suppress MiT-TFE transcription factors. Although we have focused on macrophages and microglia, modulation of lysosome degradative activity is also important in dendritic cells to ensure efficient presentation of peptide antigens^61^. Likewise, MiT-TFE overactivation may also contribute to kidney proximal tubule lysosome phenotypes, including elevated levels of multiple lysosome hydrolases, that occur in response to loss of LRRK2^20, 22, 25, 26^. Given that proximal tubule cells have a highly active endolysosomal pathway that mediates the massive uptake of proteins (including lysosomal hydrolases) from the nephron lumen in order to prevent their loss in the urine, these cells might be particularly sensitive to imbalances in lysosome homeostasis^62, 63^. More broadly, LRRK2-dependent regulation of MiT-TFE proteins may also explain previously reported effects of LRRK2 mutations and inhibitors on the levels of various lysosome proteins in additional cell types such as astrocytes and dopaminergic neurons^24, 64^.

The ability of TFE3 (and the closely related TFEB and MITF) to promote the expression of multiple genes that encode lysosomal proteins by binding to a distinct response element in their promoters has generated considerable interest in the development of strategies to enhance their activity as a strategy to boost lysosome function across multiple disease states^54, 65–68^. Beyond lysosomal targets, TFEB and TFE3 also stimulate the expression of genes related to innate immunity in macrophages^5, 69–71^. Our discovery that LRRK2 inhibition increases the expression and nuclear localization of TFE3 and TFEB suggests that LRRK2 inhibitors under development for Parkinson’s disease might have wider impacts in the treatment of other diseases associated with lysosome deficiencies and/or defects in innate immunity. Lysosome storage diseases represent an area where our data suggests new opportunities for promoting the activity of these transcription factors for therapeutic purposes. However, it remains to be determined whether the lysosome abnormalities such as those observed in the kidneys and lungs of animal models following genetic and pharmacological inhibition of LRRK2 will limit the feasibility of long term LRRK2 inhibition in humans^20, 21, 23, 25, 26, 72^. Ongoing Parkinson’s disease clinical trials will soon begin to answer these questions^36^.

An important topic for future cell biological studies will be to determine how LRRK2 fits into the complex network of regulatory machinery upstream of the MiT-TFE transcription factors^11^. The nuclear versus cytoplasmic distribution of these proteins is tightly regulated by phosphorylation by mTORC1^3^. As we did not observe any major role for LRRK2 in regulating mTORC1 activity (Fig. S9), novel targets for LRRK2 in the machinery that selectively supports mTORC1-dependent phosphorylation of MiT-TFE proteins at lysosomes may link LRRK2 kinase activity to MiT-TFE regulation^3, 11^. Alternatively, it was recently shown that itaconate, a metabolite generated in macrophages in response to innate immunity signaling, directly alkylates and activates TFEB^69^. Phosphorylation of Rab GTPases by LRRK2 could also change the composition of lysosomes by regulating intracellular membrane traffic and thus affect lysosome recruitment and phosphorylation of MiT-TFE proteins^16, 73, 74^. Recently described direct membrane remodeling properties of LRRK2 might also play a role^75^. Other signaling pathways in macrophages might intersect with MiT-TFE regulation in specific physiological contexts. For example, it was proposed that LRRK2 forms a complex with CD38 at the plasma membrane and that CD38 activation can initiate a signaling cascade that impacts lysosomes via TFEB regulation^76^. Finally, as ongoing research is still uncovering novel mechanisms of MiT-TFE regulation^77^, defining the links between LRRK2 and these transcription factors may require the discovery of new regulatory mechanisms.

In summary, we have identified a new role for LRRK2 in the transcriptional regulation of lysosomal degradative activity in macrophages and microglia via control of MiT-TFE transcription factor expression and nuclear localization. The identification of this pathway for LRRK2-dependent suppression of the degradative activity of lysosomes opens up new opportunities to define more detailed molecular mechanisms and to determine the contributions of such regulation to both normal cell biology and to Parkinson’s disease risk. Our results furthermore suggest new possibilities for exploring the use of LRRK2 inhibitors to enhance lysosome activity in other diseases associated with lysosome dysfunction.

## Methods

### Antibodies and chemicals

A summary of the antibodies used in this study can be found in Table S1. The following chemicals were used: Y-27632 Rock inhibitor (Tocris Bioscience), MLi-2 (Abcam #254528), LRRK2-IN-1 (Tocris Bioscience # 4273), EDTA (Invitrogen#15575-038).

### iPSC culture

Human female iPSCs (A18945) were purchased from Gibco. A18945 iPSCs with LRRK2 KO and G2019S mutations were kindly provided by Mark Cookson (NIH)^51^. These cells were cultured on Matrigel (Corning)-coated dishes in E8 media (Life Technologies) and passaged every 3^rd^ day using 0.5mM EDTA in phosphate buffered saline.

### Differentiation of iPSCs into Macrophages and Microglia

For microglia or macrophage differentiation, iPSCs were first differentiated to hematopoietic progenitor cells (HPSCs) using the STEMdiff kit (STEMCELL Technologies #5310) as per the manufacturer’s instructions. These cells were further differentiated into macrophages by culturing in media containing RPMI (Gibco #11875-135), 20% FBS (Gibco) and 100ng/ml MCSF (Peprotech#300-25) for 7 days via a previously published protocol^50^. HPSCs were also differentiated to microglia via an established protocol^59^.

### iPSC neuronal differentiation

Human iPSCs (WTC11 line) were differentiated for 10 days into cortical i^3^Neurons according to a previously described protocol based on the doxycycline inducible expression of Ngn2^78^.

### Bone Marrow Derived Macrophage (BMDM) differentiation and maintenance

For mouse bone marrow-derived macrophage (BMDM) primary cultures, each experiment involved age and sex matched C57Bl/6 mice between 3-8 months of age. *Lrrk2* KO (Lrrk2^tm^^1.1Mjff^) and G2019S (*Lrrk2^tm^*^1^*^.1Hlme^*) homozygous knockin mice were obtained from The Jackson Laboratory^79^. Mice were euthanized via CO_2_ or isoflurane inhalation and cervical dislocation. Femurs were collected and cavities were flushed with 5ml ice-cold PBS and the bone marrow cells were collected by centrifugation. The resulting pellet was resuspended and differentiated for 6 days in culture media containing: DMEMF12 (Gibco, #11330-032) supplemented with 20% FBS (Gibco, #16140-071), 20% L929 conditioned media, 1% penicillin-streptomycin (Gibco, # 15140-122) and 1% GlutaMAX^TM^ (Gibco, # 35050061).

### Microscopy

All microscopy experiments were performed on a Zeiss 880 Airyscan confocal microscope using a 63X plan-apochromat objective (1.46 NA). Images were acquired with Zeiss Zen Black software (RRID:SCR_018163). Further analysis was performed by using FIJI/ImageJ (RRID:SCR_003070)^80^.

### DQ-BSA assay

100,000 macrophages or microglia were seeded on Mattek glass bottom dishes. The next day cells were treated with 50 nM MLi2 or 250 nM LRRK2-IN-1 for the indicated durations where 10ug/ml DQ-BSA (Thermo Fisher Scientific, #D12051) and 50ug/ml Alexa-488 BSA (Thermo Fisher Scientific #A13100) were added during the final hour. The cells were then washed 3X with media and imaged. For DQ-BSA quantification, maximal projection images from z-stacks spanning complete cells were segmented in FIJI/ImageJ using the find maxima function. Then duplicated images were threshold by default algorithm. The segmented and thresholded images were next combined by AND function to create a mask. Finally, mean gray values were obtained by applying analyze particle function to the mask and redirecting this whole analysis to the original images.

### Immunofluorescence analysis

50,000 cells were plated on 12 mm glass coverslips in a 24 well dish. The next day cells were fixed with 4% paraformaldehyde in 0.1M phosphate buffer (pH 7.2) for 20 minutes, washed and then blocked in 5% BSA + 0.1% saponin (Sigma) in PBS for 1 hour. This was followed by overnight staining with primary antibodies at 4 degrees C, washing with PBS + 0.1% saponin and 1 hour staining with Alexa dye conjugated secondary antibodies (Thermo Fisher Scientific) at room temperature. Coverslips were then mounted on glass slides with Prolong Gold mounting media containing DAPI (Thermo Fisher Scientific).

### Quantifying the abundance of LAMP1 positive puncta

Following immunofluorescence staining and image acquisition, confocal Z-stacks (4-6 μM) were analyzed in ImageJ/Fiji. Regions of interest were defined using the “Polygon Tool” to define cell edges followed by thresholding across z-stacks. The “Analyze particle” was then used to determine the mean number of LAMP1 positive puncta per cell.

### Immunoblotting

After day 19 (human iPSC-derived macrophages) or day 40 (human iPSC-derived microglia) or day 7 (BMDM) of differentiation, cells were washed with ice-cold phosphate buffered saline and then lysed in 50mM Tris pH 7.4, 150 mM Nacl, 1% TX-100, 0.5% deoxycholate, 0.1% SDS) containing protease inhibitor and phosphatase inhibitor cocktails (Roche cOmplete and PhosSTOP) and spun at 13,000 × g for 5 min. The supernatant was collected and incubated at 95°C for 3 min in SDS sample buffer before separation by SDS–PAGE on 4–15% gradient Mini-PROTEAN TGX precast polyacrylamide gels and transferred to nitrocellulose membranes (Bio-Rad, Hercules CA). Membranes were next blocked with 5% milk in TBS with 0.1% Tween 20 and incubated with primary antibodies and then HRP-coupled secondary antibodies in 5% milk or bovine serum albumin in TBS with 0.1% Tween 20. Chemiluminescence detection was performed on a Chemidoc MP imaging station (Bio-Rad), and FIJI/ImageJ was used to quantify band intensities^80^.

### RNA isolation and qRT-PCR

Total RNA was extracted with the RNeasy kit (QIAGEN). cDNA was prepared from 1000 ng of RNA using the iScript cDNA synthesis kit (BIO-RAD). The total cDNA obtained was diluted 1:10 and 2 µl was used for qRT-PCR by using gene specific primer and SYBR green PCR mix (BIO-RAD) on BIO-RAD CFX96 real time PCR machine. Analysis was performed by calculating Δ and 2^ΔCT^ values and mRNA expression levels for genes of interest was normalized to GAPDH and represented relative to control. Oligonucleotide primer sequences are provided in Table S2.

### Nuclear/cytoplasmic fractionation

500,000 BMDMs were treated with 0.1% DMSO +/− 50 nM MLi-2 for 6 hours. After the respective treatments, cells were washed 3X in PBS, harvested in 500 μl of ice-cold hypotonic buffer [10 mM Hepes (pH 7.9), 10 mM KCl, 0.1 mM EDTA, 0.1 mM EGTA, 1 mM dithiothreitol (DTT), 0.15% NP-40], and homogenized with 20 strokes of a Dounce homogenizer. To 100μl of this homogenate, SDS (1% final) and 25 U of Benzonase (Novagen) were added, and the rest of the homogenate was spun at 4°C for 5 min at 14,000 rpm. The supernatant (cytoplasmic fraction) was collected into a new tube, and the pellet (nuclear fraction) was resuspended in 200 μl of high-salt buffer (20 mM Hepes, 400 mM NaCl, 1 mM EDTA, 1 mM EGTA, 1 mM DTT, 0.5% NP-40) and solubilized with SDS (1% final) in the presence of 25 U of Benzonase. The protein concentration was measured with the BCA reagent (Thermo Scientific), and samples were subsequently analyzed by immunoblotting.

### siRNA-mediated knockdown of TFE3

100,000 cells were plated in a 100mm cell culture dish. The next day, siRNA transfections were performed with 20 μM siRNA, 40 μl RNAiMAX transfection reagent (Invitrogen), and 1,000 μl Opti-MEM (Invitrogen) that was added to 8 ml of cell culture media. Experiments were performed 2 days post transfection. TFE3 (5′-CAAACAGUACUUGUCUAC-3’ and 5’-AAGUGUGGUAGACAAGU-3) and control (5’-AUACGCGUAUUAUACGCGAUUAACGAC-3’ and 5’-CGUUAAUCGCGUAUAAUACGCGUAT - 3’) siRNAs were purchased from Integrated DNA. The TFE3 siRNA has 9 and 4 mismatches respectively with the corresponding sequences from TFEB and MITF.

### Statistical analysis

Statistical analysis was performed using Graphpad Prism 8 software (RRID:SCR_002798). Detailed statistical information (specific test performed, number of independent experiments, and p values) is presented in the respective figure legends.

## Acknowledgements

We are grateful for microscopy resources provided by the Yale Program in Cellular Neuroscience, Neurodegeneration and Repair imaging facility. This research was supported by grants from the Parkinson’s Foundation, the Ludwig Foundation and Aligning Science Across Parkinson’s disease (ASAP-000580) through the Michael J. Fox Foundation for Parkinson’s Research (MJFF). Agnes Roczniak-Ferguson (Yale), Amanda Bentley-DeSousa (Yale), Francesca Filippini (Yale), Timothy Ryan (Cornell), Berrak Ugur (Yale) and Pietro De Camilli (Yale) provided valuable feedback. We thank Mark Cookson (NIH) for sharing LRRK2 KO and G2019S knockin human iPSCs. We appreciate assistance from Maria Lara-Tejero (Yale) and Jorge Galan (Yale) with LRRK2 mutant mice. The authors do not have any conflicts to declare.

## Author contributions

NY and SMF designed experiments. NY performed all experiments. NY and SMF analyzed data and prepared the manuscript.

## Supplemental Material

Fig. S1 shows the impact of LRRK2 perturbations on lysosomal protease activity in mouse BMDMs. Fig. S2 shows the impact of LRRK2 inhibition and KO on lysosome protein levels in mouse BMDMs. Fig. S3 shows the impact of LRRK2 inhibition and the G2019S mutation on lysosome protein levels in mouse BMDMs. Fig. S4 shows that LRRK2 KO and MLi2 treatment increase the abundance of both immature and mature forms multiple lysosome hydrolases in human iPSC derived macrophages. Fig. S5 presents the regulation of TFE3 nuclear localization by LRRK2 in mouse BMDMs. Fig. S6 shows that LRRK2 negatively regulates TFEB nuclear localization on human iPSC-derived macrophages Fig. S7 shows the lack of LRRK2-dependent regulation of TFE3 localization in human iPSC-derived neurons. Fig. S8 demonstrates that LRRK2 regulates the abundance of TFE3, TFEB and MITF in mouse BMDMs. Fig. S9 shows LRRK2 does not control mTORC1 activity in human iPSC-derived macrophages. Table S1 describes antibodies and Table S2 defines oligonucleotide primers.

## Notes

### Competing Interest Statement

The authors have declared no competing interest.

### Summary of Updates

This version has been updated to include new data and interpretation.

