## Supplemental Material for "LRRK2 Suppresses Lysosome Degradative Activity in Macrophages and Microglia Through MiT-TFE Transcription Factor Inhibition"

**This section includes:**

Figures S1 to S9
Table S1 and S2

**Supplemental Figure 1**

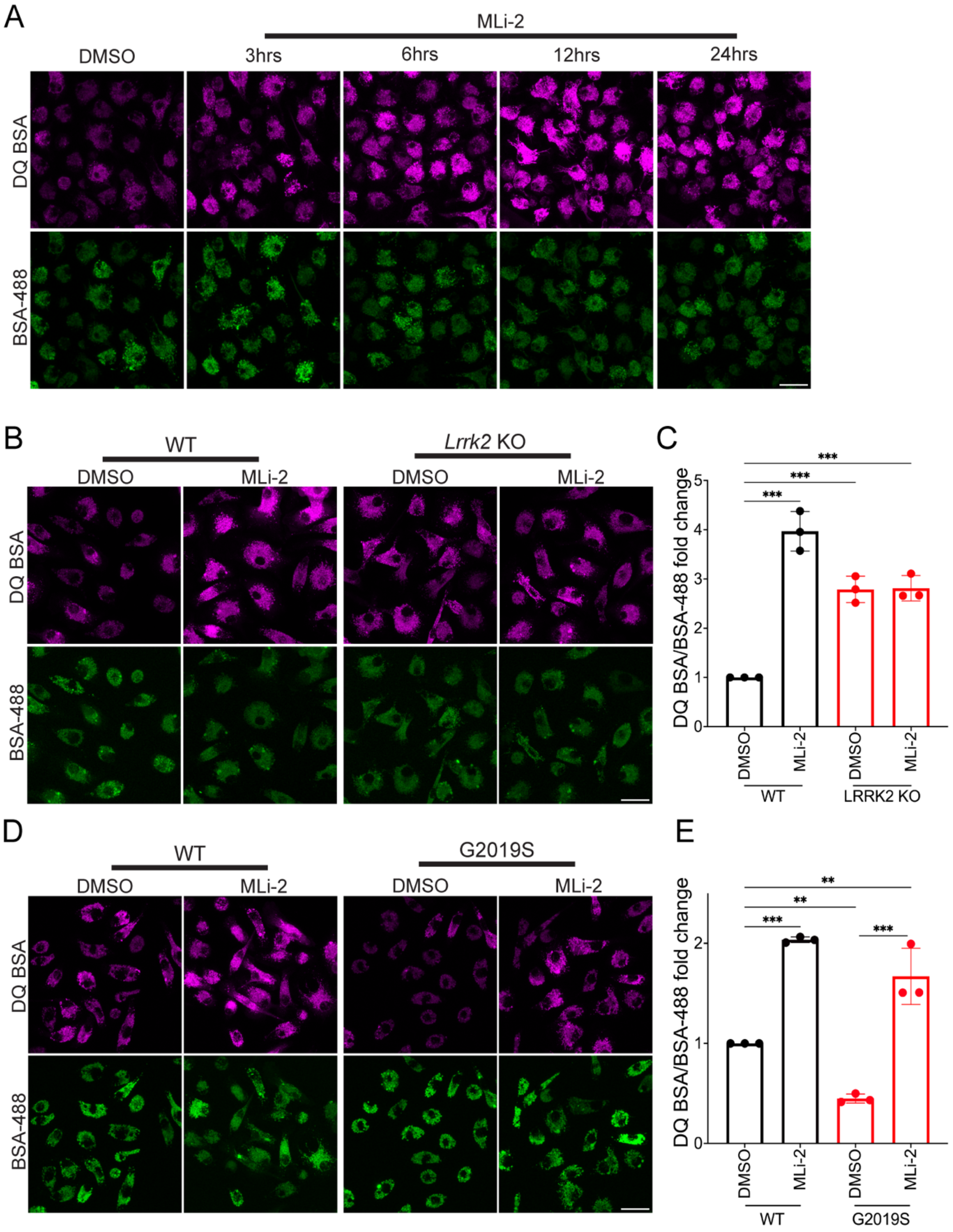

**Figure S1: LRRK2 negatively regulates lysosomal proteolytic activity in mouse bone marrow derived macrophages (BMDMs). (A)** Confocal micrographs showing the DQ-BSA and Alexa488-BSA fluorescence in WT cells treated with 0.1% DMSO or 50 nM MLi-2 in for 3,6,12 and 24 hours. Scale bar, 10 µm. **(B)** Confocal micrographs showing DQ-BSA and Alexa488-BSA fluorescence in WT and LRRK2 KO BMDMs treated for 3 hours with 0.1% DMSO or 50 nM MLi2. Scale bar, 10 µm. **(C)** Quantification of the DQ-BSA/Alexa488-BSA fluorescence. The mean fluorescence intensity of DQ-BSA was normalized to Alexa488-BSA and plotted relative to the DMSO control. The data was collected from 3 independent experiments with 60-80 cells per experiment. **(D)** Confocal micrographs showing DQ-BSA and Alexa488-BSA fluorescence in WT and LRRK2 G2019S knockin mouse BMDMs that were treated with 0.1% DMSO or 50 nM MLi2 for 3 hours. Scale bar, 10 µm. **(E)** Quantification of DQ-BSA/Alexa488-BSA fluorescence in WT and LRRK2 G2019S macrophages. The data was collected from 3 independent experiments with 60-80 cells per experiment (error bars show mean ± SEM, one-way ANOVA, * $p<0.05$, ** $p< 0.01$, ***$p< 0.001$).

**Supplemental Figure 2**

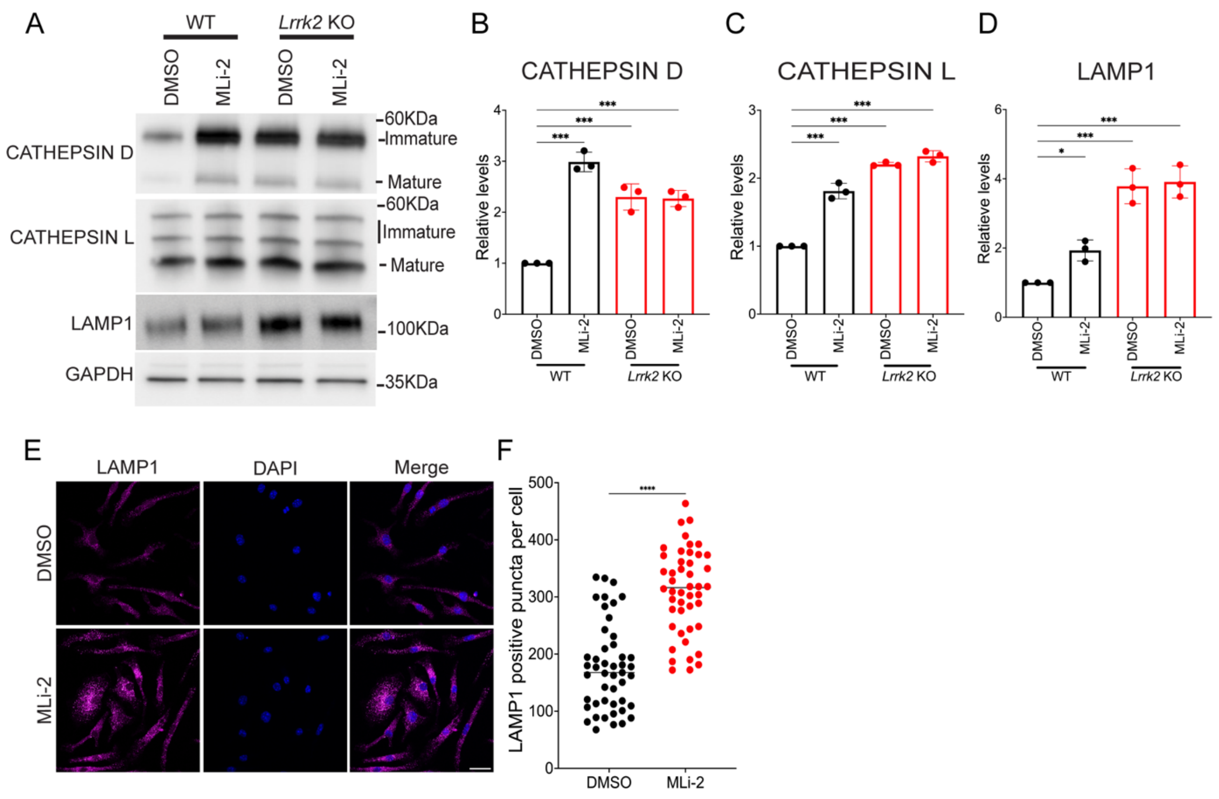


**Figure S2: LRRK2 KO and MLi2 treatment increase the abundance of multiple lysosome proteins in mouse bone marrow derived macrophages. (A)** Immunoblotting analysis of WT and LRRK2 KO mouse BMDMs treated with 0.1% DMSO or 50 nM MLi-2 for 6 hours. **(B-D)** Quantification of immunoblots in A. Data was plotted in relative to the DMSO-treated WT cells (n=3 independent experiments). **(E)** Confocal micrographs showing the LAMP1 immunostaining in mouse BMDMs treated with 0.1% DMSO or 50 nM MLi-2 for 6 hours. Scale bar, 10 μm. **(F)** Quantification of LAMP1 positive puncta shown in E (n=3 experiments with 17-20 cells analyzed per experiment. For all panels error bars show mean ± SEM, one-way ANOVA, * p<0.05, ** p< 0.01, *** p< 0.001).

**Supplemental Figure 3**

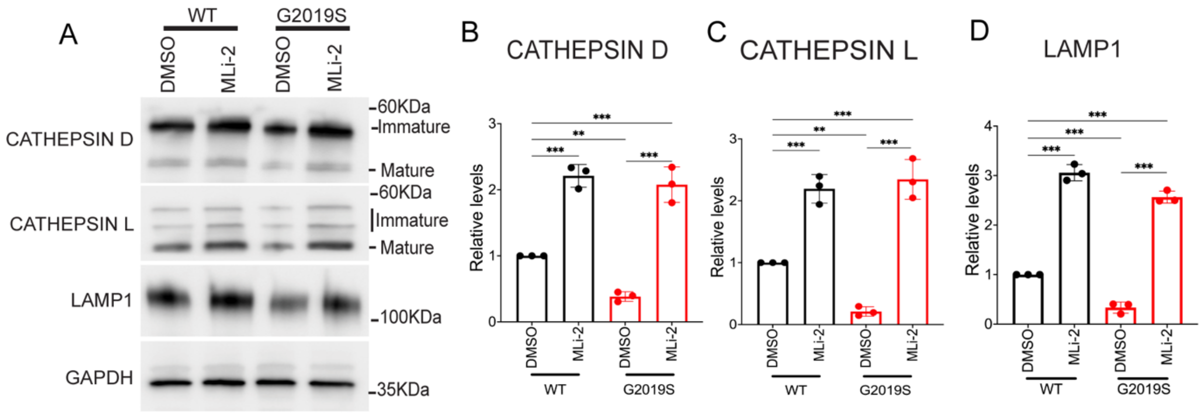


**Figure S3: LRRK2 G2019S knockin mouse bone marrow derived macrophages have reduced levels of multiple lysosome proteins. (A)** Immunoblot analysis of WT and LRRK2 G2019S mutant BMDM's treated with 0.1% DMSO or 50 nM MLi-2 for 6 hours. **(B-D)** Quantification of immunoblots from panel A. Data was plotted relative to the DMSO-treated WT cells (n=3, error bars show mean ± SEM, one-way ANOVA, * $p<0.05$, ** $p< 0.01$, *** $p< 0.001$).

**Supplemental Figure 4**

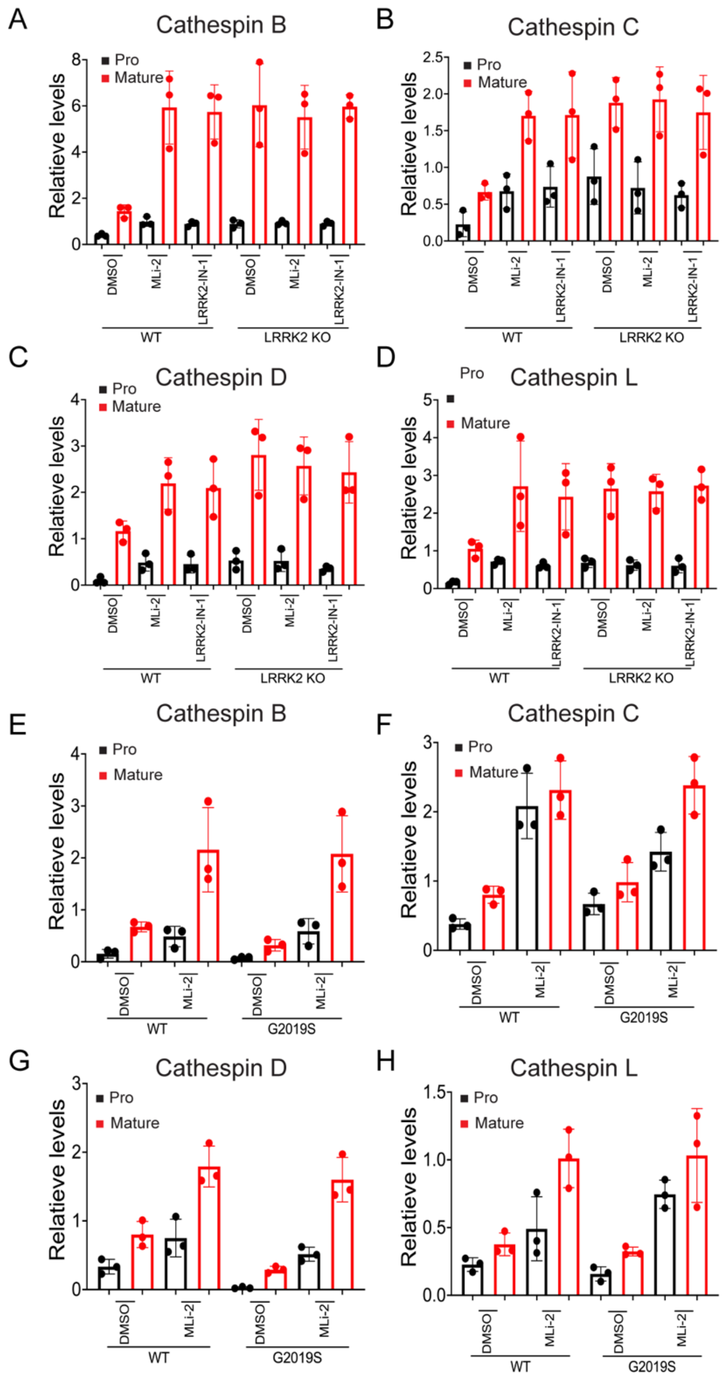


**Figure S4: LRRK2 KO and MLi2 treatment increase the abundance of multiple lysosome proteins in human iPSC derived macrophages. (A-D)** Bar graphs showing the quantification of immature and mature cathepsin B, C, D and L levels in LRRK2 KO and MLi-2 treated conditions. This analysis was performed on the data presented in Figure 2. **(E-H)** Bar graphs showing the quantification of immature and mature cathepsin B, C, D and L levels in G2019S knockin and MLi-2 treated conditions (based on re-analysis of results from Figure 3).

### Supplemental Figure 5

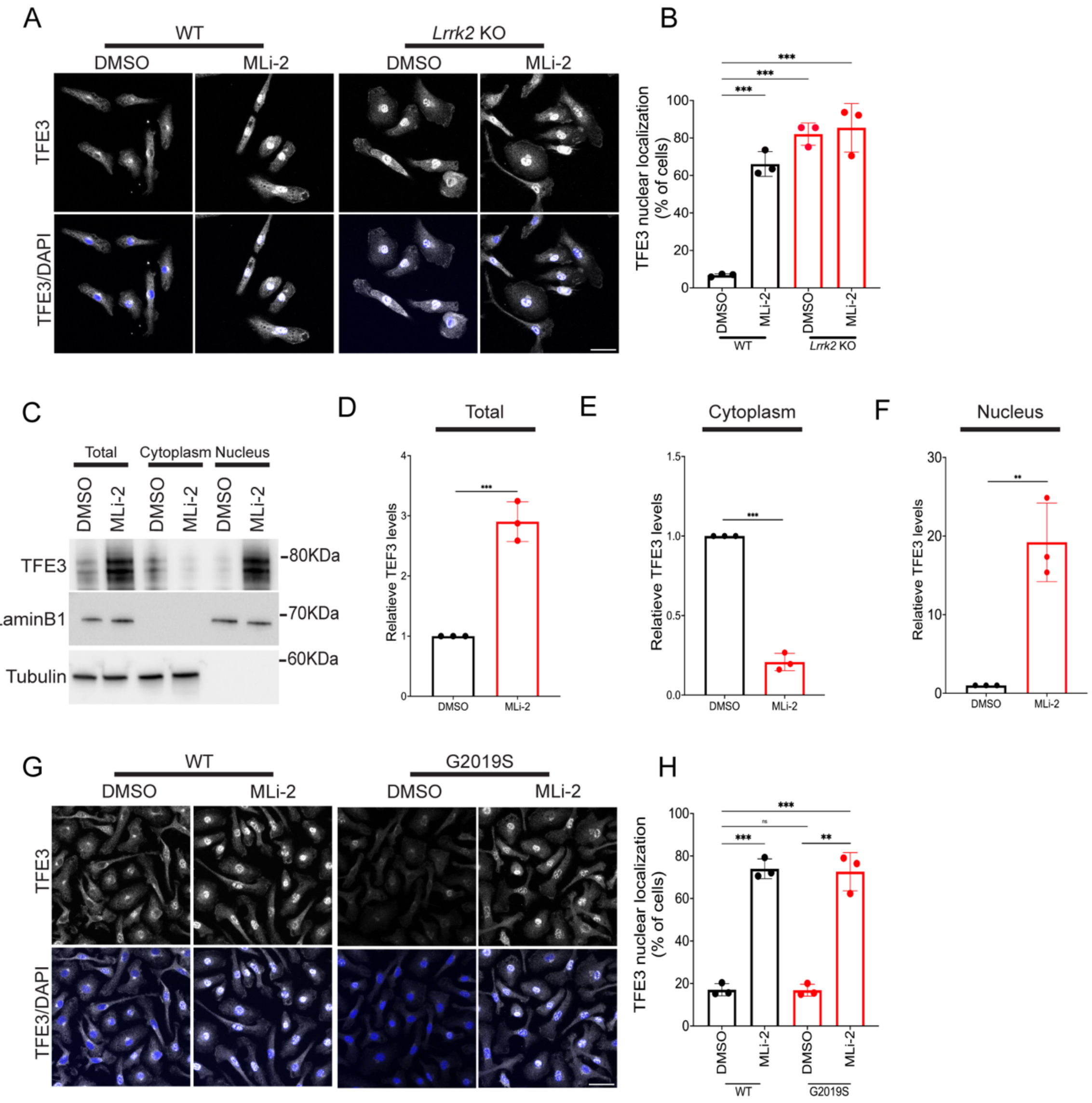


**Figure S5: LRRK2 negatively regulates the nuclear localization of TFE3 in mouse bone marrow derived macrophages. (A)** Immunofluorescence confocal micrographs showing the sub-cellular localization of TFE3 in WT and LRRK2 KO

BMDMs that were treated with 0.1% DMSO or 50 nM MLi-2 for 3 hours. Scale bar, 10 µm. **(B)** Quantification of the % of cells with nuclear>cytoplasmic TFE3. Data was collected from 3 independent experiments with approximately 50-60 cells analyzed per experiment. **(C)** Immunoblots showing the levels of TFE3 in total cell lysate, cytoplasmic and nuclear fractions from BMDM's treated with 0.1% DMSO or 50nM MLi-2 for 6 hours. **(D-F)** Quantification of immunoblots shown in C. **(G)** Immunofluorescence confocal micrographs showing the sub-cellular localization of TFE3 in WT and LRRK2 G2019 mutant BMDMs that were treated with 0.1% DMSO or 50 nM MLi-2 for 3 hours. Scale bar, 10 µm. **(H)** Quantification of the % cells with nuclear>cytoplasmic TFE3 under the indicated conditions. Data was collected from 3 independent experiments with approximately 50-60 cells per experiment. Scale bars = 10µm, error bars show mean ± SEM, error bars show mean ± SEM, one-way ANOVA, * p<0.05, ** p< 0.01, ***p< 0.001.

**Supplemental Figure 6**

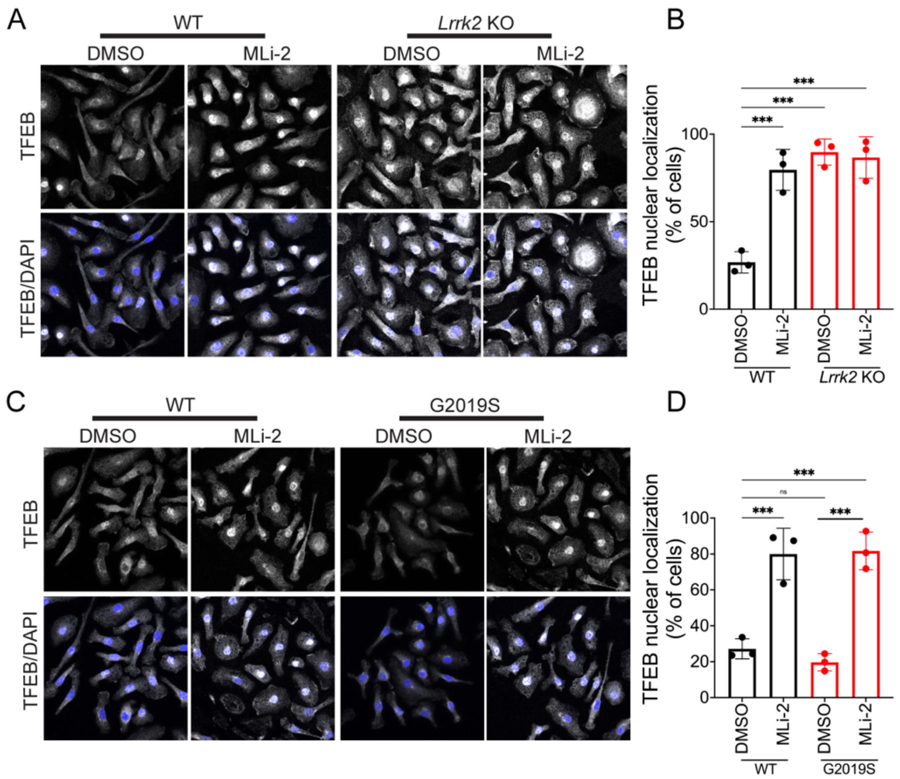

**Figure S6: LRRK2 negatively regulates the nuclear localization of TFEB in mouse bone marrow derived macrophages. (A)** Immunofluorescence, confocal micrographs showing the sub-cellular localization of TFEB in WT and LRRK2 KO BMDMs that were treated with 0.1% DMSO or 50 nM MLi-2 for 3 hours. Scale bar, 10 µm. **(B)** Quantification of the % of cells with nuclear>cytoplasmic TFEB. Data was collected from 3 independent experiments with approximately 50-60 cells analyzed per experiment. **(C)** Immunofluorescence, confocal micrographs showing the sub-cellular localization of TFEB in WT and LRRK2 G2019 mutant BMDMs that were treated with 0.1% DMSO or 50 nM MLi-2 for 3 hours. Scale bar, 10 µm. **(D)** Quantification of the % cells with nuclear>cytoplasmic TFEB under the indicated conditions. Data was collected from 3 independent experiments with approximately 50-60 cells per experiment. Scale bars = 10µm, error bars show mean ± SEM, one-way ANOVA, * $p<0.05$, ** $p< 0.01$, ***$p< 0.001$.

**Supplemental Figure 7**

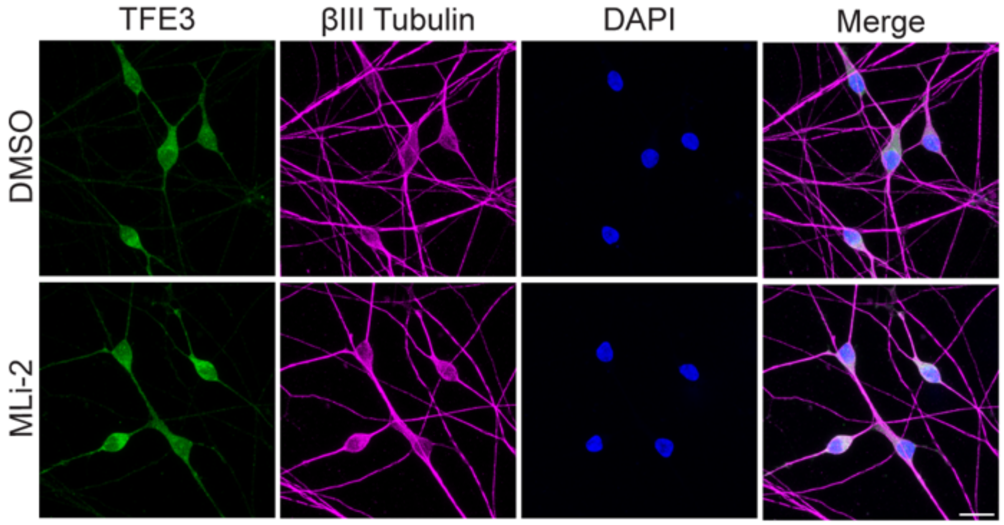


**Figure S7.** Confocal micrographs showing TFE3 localization in human iPSC derived cortical neurons treated with 0.1% DMSO or 50nM MLi-2 for 6 hours. Scale bar = 10 µm.

**Supplemental Figure 8**

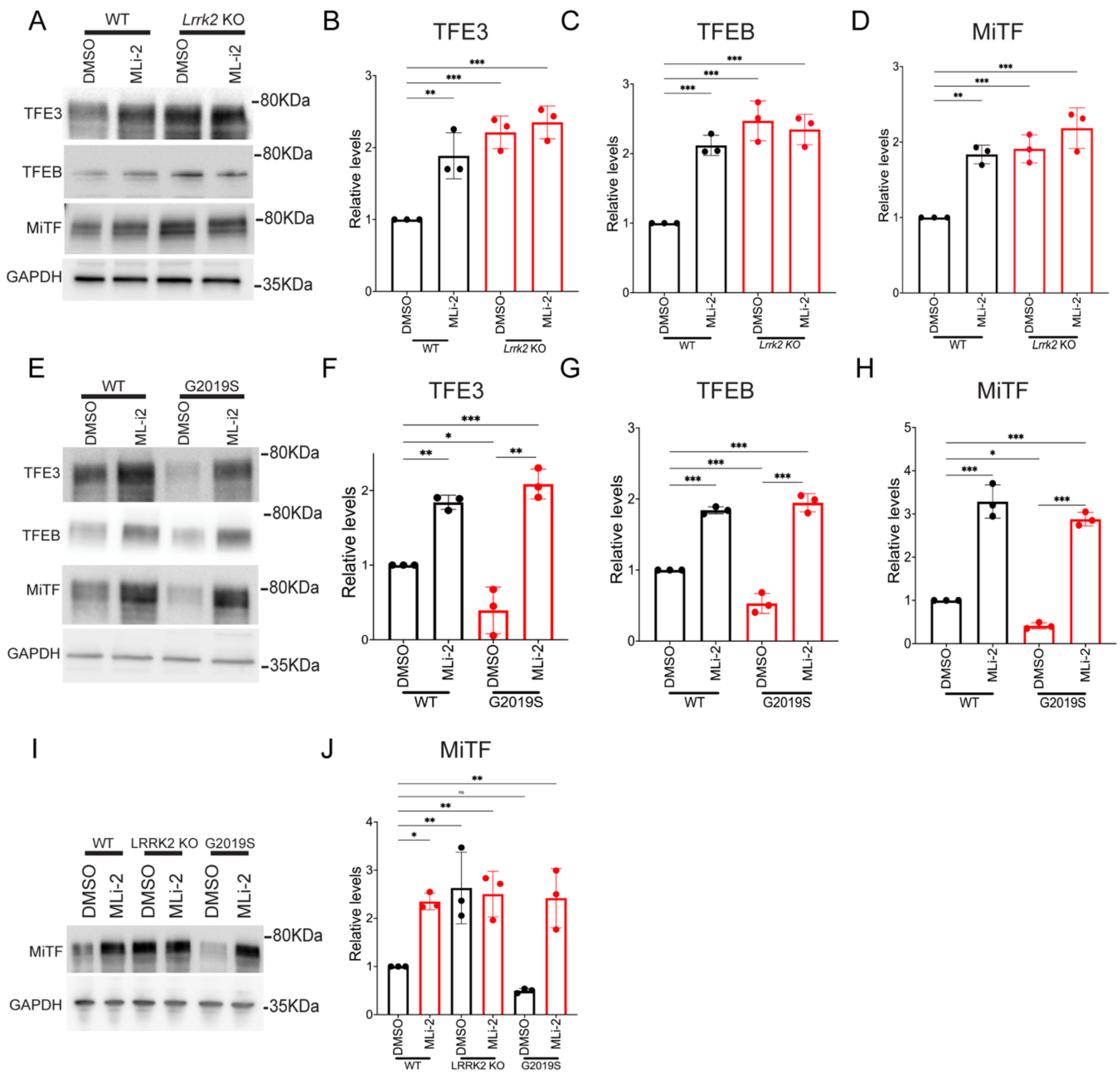


**Figure S8: LRRK2 negatively regulates the abundance of TFE3, TFEB and MITF in mouse bone marrow derived macrophages. (A)** Immunoblots showing the levels of TFE3, TFEB and MiTF in WT and LRRK2 KO cells treated with 0.1% DMSO or 50nM MLi-2 for 6 hours. **(B-D)** Quantification of immunoblots from panel A. Data was collected from 3 independent experiments, normalized to GAPDH and plotted relative to DMSO-treated WT cells. **(E)** Immunoblots showing the levels of TFE3, TFEB and MITF in WT and LRRK2 G2019S macrophages. **(F-H)** Quantification of immunoblots in panel E. Data was collected from 3 independent experiments, normalized to GAPDH and represented relative to the DMSO-treated

WT macrophages. **(I)** Immunoblots showing the levels of MITF in WT, LRRK2 KO and LRRK2 G2019S BMDMs cells treated with 0.1% DMSO or 50nM MLi-2 for 6 hours. **(J)** Quantification of immunoblots shown in M. Data was collected from 3 independent experiments, normalized to GAPDH and represented relative to the control siRNA treated WT macrophages. Error bars show mean ± SEM, one-way ANOVA, * $p<0.05$, ** $p< 0.01$, *** $p< 0.001$.

**Supplemental Figure 9**

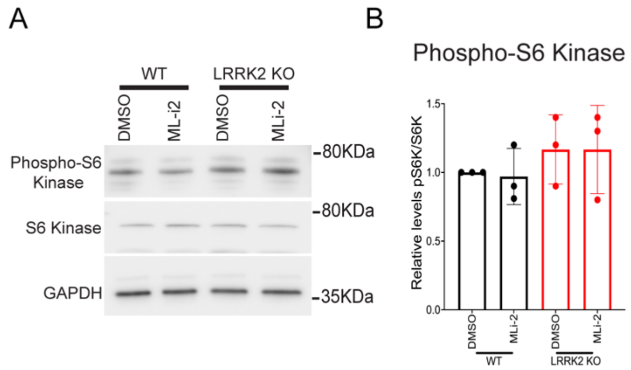


**Figure S9: LRRK2 inhibition does not affect the mTORC1-dependent phosphorylation of ribosomal protein S6 kinase. (A)** Immunoblots from human iPSC-derived macrophages showing the levels of phosphorylated S6 Kinase (T389), total S6 kinase and GAPDH in WT and LRRK2 KO cells treated with 0.1% DMSO or 50nM MLi-2 for 6 hours. **(B)** Quantification of immunoblots where data was collected from 3 independent experiments, normalized to GAPDH and plotted relative to DMSO-treated WT cells. Error bars show mean ± SEM.

**Table S1: Sources of antibodies used.**

| Antibody | Company | Catalog Number | RRID |
|---|---|---|---|
| Human cathepsin B | R&D Systems | AF953 | AB_355738 |
| Human cathepsin C | R&D Systems | AF1071 | AB_2086965 |
| Human Cathepsin D | R&D Systems | AF1014 | AB_2087218 |
| Mouse Cathepsin D | R&D Systems | AF1029 | AB_2087094 |
| Human cathepsin L | R&D Systems | AF952 | AB_355737 |
| Mouse cathepsin L | R&D Systems | AF1515 | AB_2087690 |
| Human GBA | R&D Systems | MAB7410 | AB_2938856 |
| GAPDH | EnCor Biotechnology Inc | MCA-1D4 | AB_2107599 |
| Human LAMP1 | Cell signaling Technology | 9091 | AB_2687579 |
| Mouse LAMP1 | DSHB | 1D4B | AB_2134500 |
| MITF | Cell signaling Technology | 12590 | AB_2616024 |
| MITF | Abcam | Ab303530 | AB_2938857 |
| S6K | Cell Signaling Technology | 9202 | AB_331676 |
| S6K-Phospho-T389 | Cell Signaling Technology | 9205 | AB_330944 |
| Human TFEB | Cell Signaling Technology | 37785 | AB_2799119 |
| Mouse TFEB | Proteintech | 13372-1-AP | AB_2199611 |
| TFE3 | Sigma | HPA023881 | AB_1857931 |

**Table S2. Oligonucleotide primer sequences used for qRT-PCR assays.**

| | |
|---|---|
| CTSL-FP | CTGGTGGTTGGCTACGGATT |
| CTSL-RP | CTCCGGTCTTTGGCCATCTT |
| CTSD-FP | AACTGCTGGACATCGCTTGCT |
| CTSD-RP | CATTCTTCACGTAGGTGCTGGA |
| TFE3-FP | ACTACTGTCAGCAACTCCTG |
| TFE3-RP | CTGTCGTTAACGTTGAATCGCC |
| GAPDH-FP | TGCACCACCAACTGCTTAGC |
| GAPDH-RP | GGCTAGGACTGTGGTCATGAG |
| LAMP1-FP | CCTGGGTGCCACTAACACAT |
| LAMP1-RP | CACACTTTTCCCCATAGCG |
| GBA-FP | ATGCAAGAGTGAATGGGAAGG |
| GBA-RP | TTCATTCTCCGCTGTCACTC |
